# Modular Cell Line for Scalable and Rapid In Vitro Evaluation of Chimeric Antigen Receptors

**DOI:** 10.1101/2024.08.10.604092

**Authors:** Simon Dubovik, Dmitri Dormeshkin, Alexandr Migas

## Abstract

The development of novel Chimeric Antigen Receptor T-cell (CAR-T-cell) products relies on continuous *in vitro* testing of new antigen/antibody pairs, as well as modification of CAR composition itself. Since any Tumor-Associated Antigens (TAA) have been chosen and proven to be specific, and before the first *in vivo* studies these CAR compositions have to be extensively validated *in vitro*. Conventional *in vitro* tests rely on TAA expressing immortalized cell lines or immortalized cell lines derivatives engineered to stably express antigen of interest, making this type of model not fully scalable when it comes to multiple TAA/antibody pairs. Therefore, there is a need for a representative, scalable model for preclinical assessment. Here, we introduce a novel model cell line approach that can be easily adapted for different experiments by anchoring ALFA-tagged proteins on the cell surface using an anti-ALFA-tag membrane single-domain antibody. This modular multi-component system has the potential to reduce novel CAR *in vitro* evaluation procedures from months to weeks.

## 1 Introduction

The simplest models are indispensable for the initial evaluation of novel Chimeric Antigen Receptor (CAR) T-cell constructs. The development of these products involves stages such as target identification, antibody selection, and CAR cassette assembly, followed by *in vitro* cytotoxic activity measurements and *in vivo* evaluation. The *in vitro* tests typically rely on a low-level models (Si et al., 2022) like Tumor Associated Antigen (TAA)-positive tumor cell lines (Lei et al., 2024; Palomba et al., 2022; Zhang et al., 2023) or TAA-modified immortalized cell lines thus providing meaningful information about CAR functionality in corresponding cells.

*In vitro* CAR-T-cell functional activity can be analyzed with different models, including antigen-coated beads/plates, primary TAA-harboring tumor cell lines and TAA knock-in established cell line (Si et al., 2022). The first one, or antigen-coated beads/plates results in CAR activation that can be evaluated by the expression of multiple molecules (CD69, CCR7, CD62L, PD-1, TIM-3, etc.) (Liu et al., 2023) as well CAR-T cell proliferation. Additionally, culture supernatants can be examined for the presence of characteristic cytokines following antigen presentation. However, it is not possible to detect direct killing events mediated by 2nd or 3rd generation CARs using antigen-coated beads/plates.

In contrast, the utilization of living targets reveals the primary function of CAR-T-cells, which is direct killing. These living targets include established cell lines and engineered cell lines. In some cases, maintenance of a particular established cell line is complicated by antigen instability during cultivation (Pandita et al., 2004). This phenomenon can be caused by altered conditions during *in vitro* culturing, leading to changes in the membrane protein repertoire and corresponding TAA content.

In some instances, multiple immortalized cell lines are available for specific antigens. This scenario is observed with CD19, where some B-cell leukemia and lymphoma derived cell lines can be utilized. These cell lines exhibit varying target antigen densities, from several thousands to several tens of thousands of CD19 molecules per cell, which is pivotal for optimal CAR-T cell functionality.

According to published data, the absolute expression values vary considerably and depend on the materials used and methods of determination (Table 1).

**Table 1.**
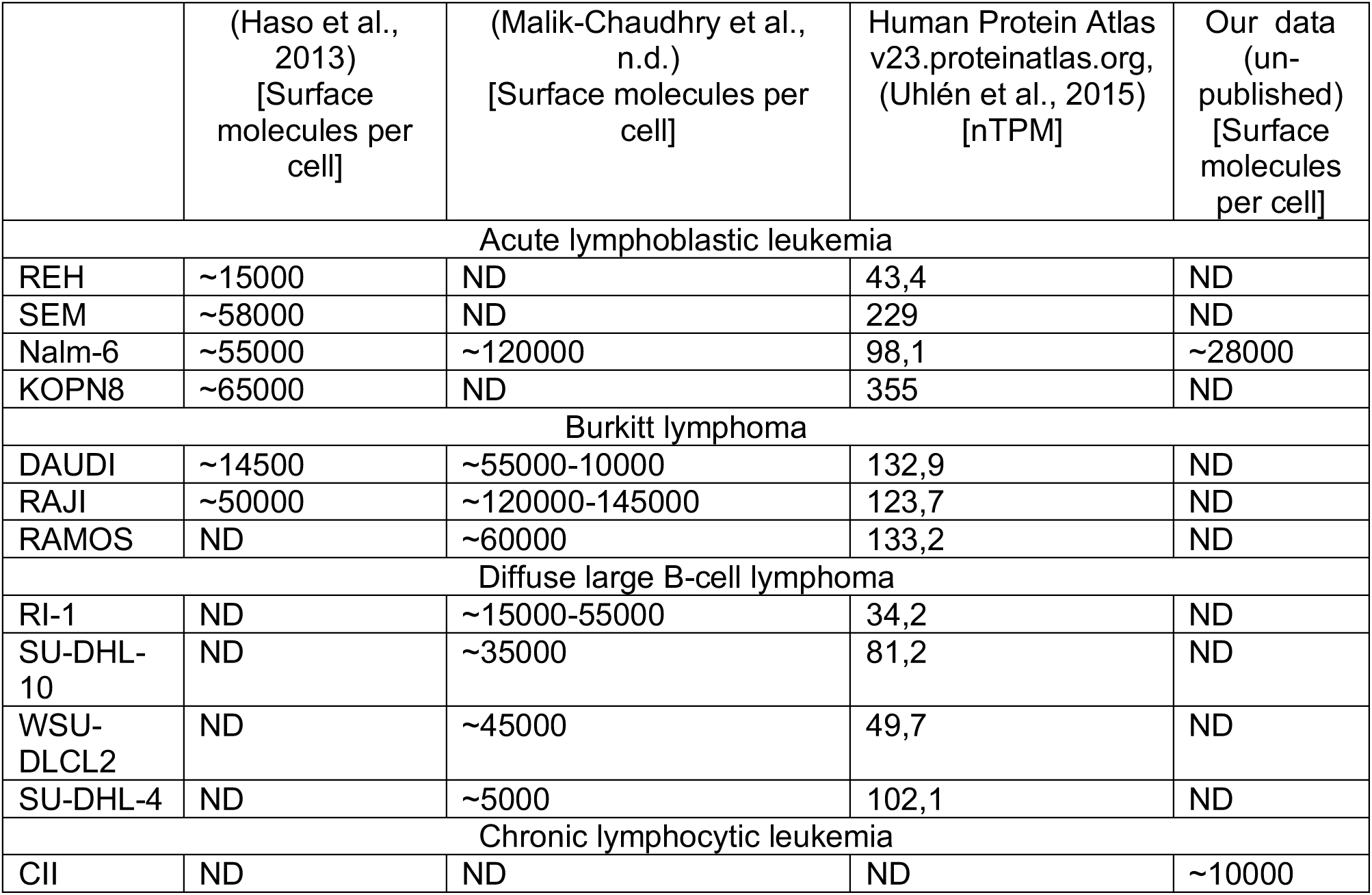
Expression of CD19 protein in various model immortalized cell lines.

|  | (Haso et al., 2013)<br>[Surface molecules per cell] | (Malik-Chaudhry et al., n.d.)<br>[Surface molecules per cell] | Human Protein Atlas v23.proteinatlas.org, (Uhlén et al., 2015)<br>[nTPM] | Our data (un-published)<br>[Surface molecules per cell] |
| --- | --- | --- | --- | --- |
| **Acute lymphoblastic leukemia** |  |  |  |  |
| REH | ~15000 | ND | 43,4 | ND |
| SEM | ~58000 | ND | 229 | ND |
| Nalm-6 | ~55000 | ~120000 | 98,1 | ~28000 |
| KOPN8 | ~65000 | ND | 355 | ND |
| **Burkitt lymphoma** |  |  |  |  |
| DAUDI | ~14500 | ~55000-10000 | 132,9 | ND |
| RAJI | ~50000 | ~120000-145000 | 123,7 | ND |
| RAMOS | ND | ~60000 | 133,2 | ND |
| **Diffuse large B-cell lymphoma** |  |  |  |  |
| RI-1 | ND | ~15000-55000 | 34,2 | ND |
| SU-DHL-10 | ND | ~35000 | 81,2 | ND |
| WSU-DLCL2 | ND | ~45000 | 49,7 | ND |
| SU-DHL-4 | ND | ~5000 | 102,1 | ND |
| **Chronic lymphocytic leukemia** |  |  |  |  |
| CII | ND | ND | ND | ~10000 |

Furthermore, it is essential to incorporate a negative control cell line during experiments to confirm the absence of nonspecific cytotoxicity. Since CAR-T function does not rely on a highly-sensitive TCR capable of recognizing a single antigen (Huang et al., 2013; Irvine et al., 2002) at the MHC complex, but instead engages numerous CAR molecules by clustering (Mukherjee et al., 2017), this process is likely dependent on accessible antigen density during recognition by CAR, while the threshold of activation can vary depending on the CAR construct itself (Fujiwara et al., 2020). Interestingly, according to the Super-resolution microscopy the cells bearing myeloma cells that express less than 1350 CD19 molecules are classified as CD19neg according to Flow Cytometry. In addition it was shown that less than 100 CD19 molecules is sufficient to elicit antitumor activity of CAR T cell (Nerreter et al., 2019). But it is still not clear whether new antigen-antibody pair would be as successful as it was in the case of anti-CD19 FMC63-based CAR which resulted in ∼40% 2-year progression-free survival (Schuster et al., 2022). This is the reason why CAR evaluation should include experiments on cell cultures with varying antigen densities.

Notably, the use of different cell lines might introduce additional influences from other surface molecules exhibited by different cell types. For example, ICAM-1 facilitates the formation of a productive synapse by interacting with LFA-1 on T-cells. According to the Human Protein Atlas, the expression levels of ICAM-1 on Daudi and KM-H2 differ by a factor of 10 (34 and 387 nTPM, respectively), indicating a wide expression range of this molecule. It was reported that the expression of ICAM-1 correlates with the activation and killing efficacy of CD20 CAR (Kantari-Mimoun et al., 2021) against Raji cells However, another study shows a redundant role of LFA-1 in the case of HER2-expressing MC57 target cell recognition by the corresponding CAR (Davenport et al., 2018). It is not clear whether any particular CAR will be affected by varying LFA-1 interactions on different cell lines.

To mitigate the influence of these variables on experimental outcomes, using a single cell line as a negative control alongside antigen-positive targets post-genetic modification is advantageous. This experimental strategy allows for the specific evaluation of CAR functionality independently of other factors, facilitating direct comparisons among different targets with the cognate antigen-recognition domains of CAR. In this context, the K-562 cell line emerges as a promising candidate due to its inherent susceptibility to apoptosis, ease of handling, and capability for efficient genetic modification.

Despite the advantages provided by the usage of engineered cells for in vitro CAR evaluation, challenges have emerged with the utilization of numerous cell lines. The cloning, modification, selection, and characterization of every target line are highly time-consuming and labor-intensive processes that usually are not scalable for high-throughput pipelines. Even more complicated it becomes with the introduction of numerous co-stimulatory molecules as well as novel antibody allele variant preference determination.

Here we present a novel model cell line that can be easily modified by anchoring any tagged protein on the cell surface using an anti-tag single-domain antibody (VHH) on the cell surface. The antibody–tag system is based on the ALFA-tag – VHH interaction presented by Götzke and colleagues (Götzke et al., 2019). This multi-component system can reduce *in vitro* CAR evaluation procedures from months to weeks by transient anchoring of the target antigen on the cell surface avoiding a highly time-consuming procedure of lentiviral transduction or other widespread methods of stable genetic modification for every particular antigen (Figure 1). Together with phage display techniques for antibody selection, it is possible to optimize antibody validation after the selection step directly using the same antigen as for the selection. That allows us to perform mass *in vitro* testing of different antigens and their cognate antibodies for further CAR T-cell production.

**Figure 1.**
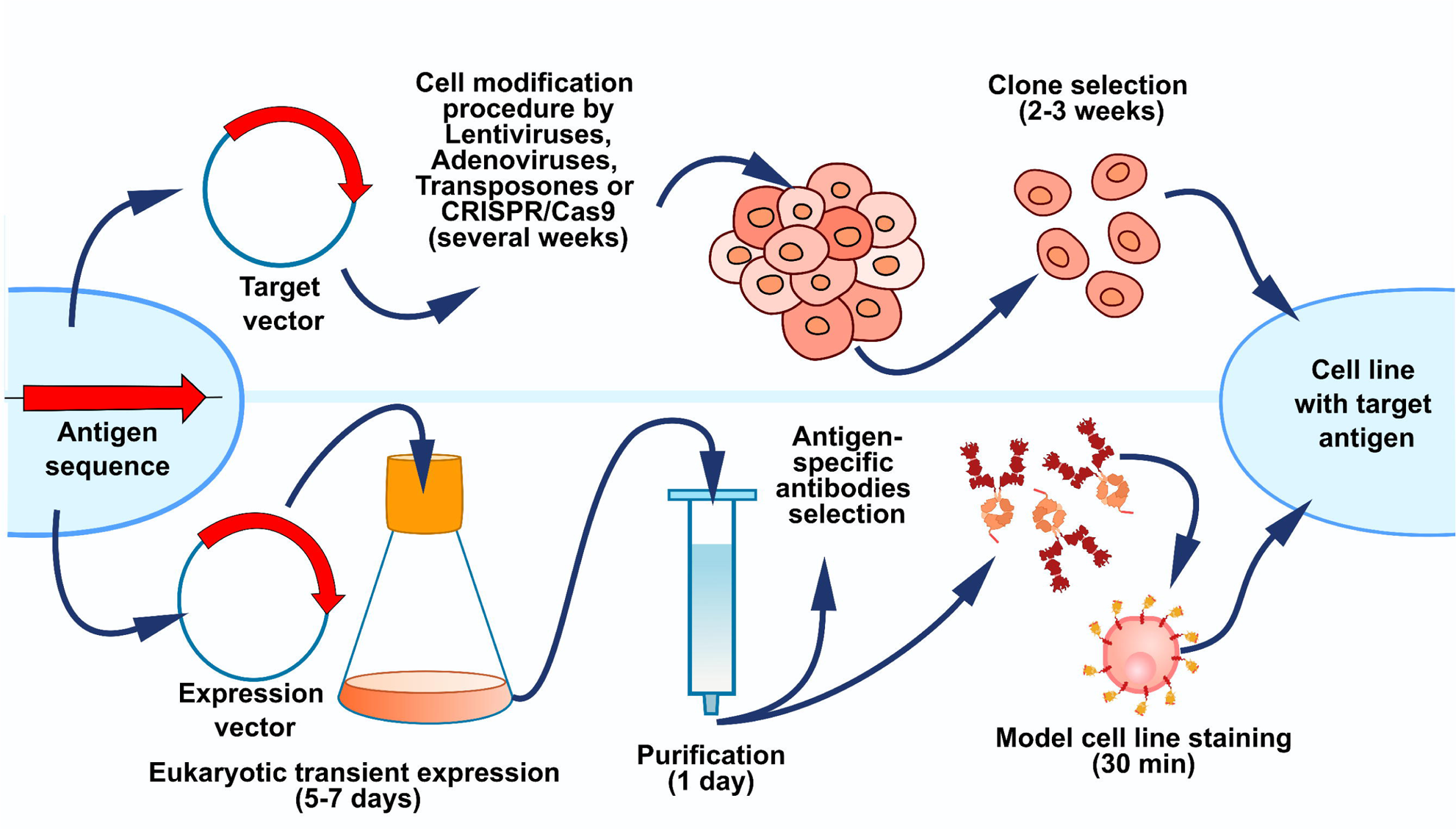
Schematic comparison of approaches. The upper section represents the classical TAA knock-in cell line and the lower – the proposed solution.

## 2 Results

### 2.1 Assembly and verification of the lead constructs

Initially, the interaction between anti-ALHA-tag VHH (aNab) and ALFA-tagged protein of interest was assessed utilizing the Octet R2 system (Sartorius, Germany). This involved the expression of Аvi-tagged aNab (Supplementary Figure 1a) and ALFA-tagged CD19 protein. The aNab:ALFA-tag interaction was registered by biosensor with immobilized Streptavidin. The testing procedure included aNab∼Аvi-tag immobilization followed by its interaction with ALFA-taged CD19 in different concentrations, resulting in binding and dissociation curves. Based on Octet R2 analysis, the estimated dissociation constant was approximately 0.5 nM (Figure 2).

**Figure 2.**
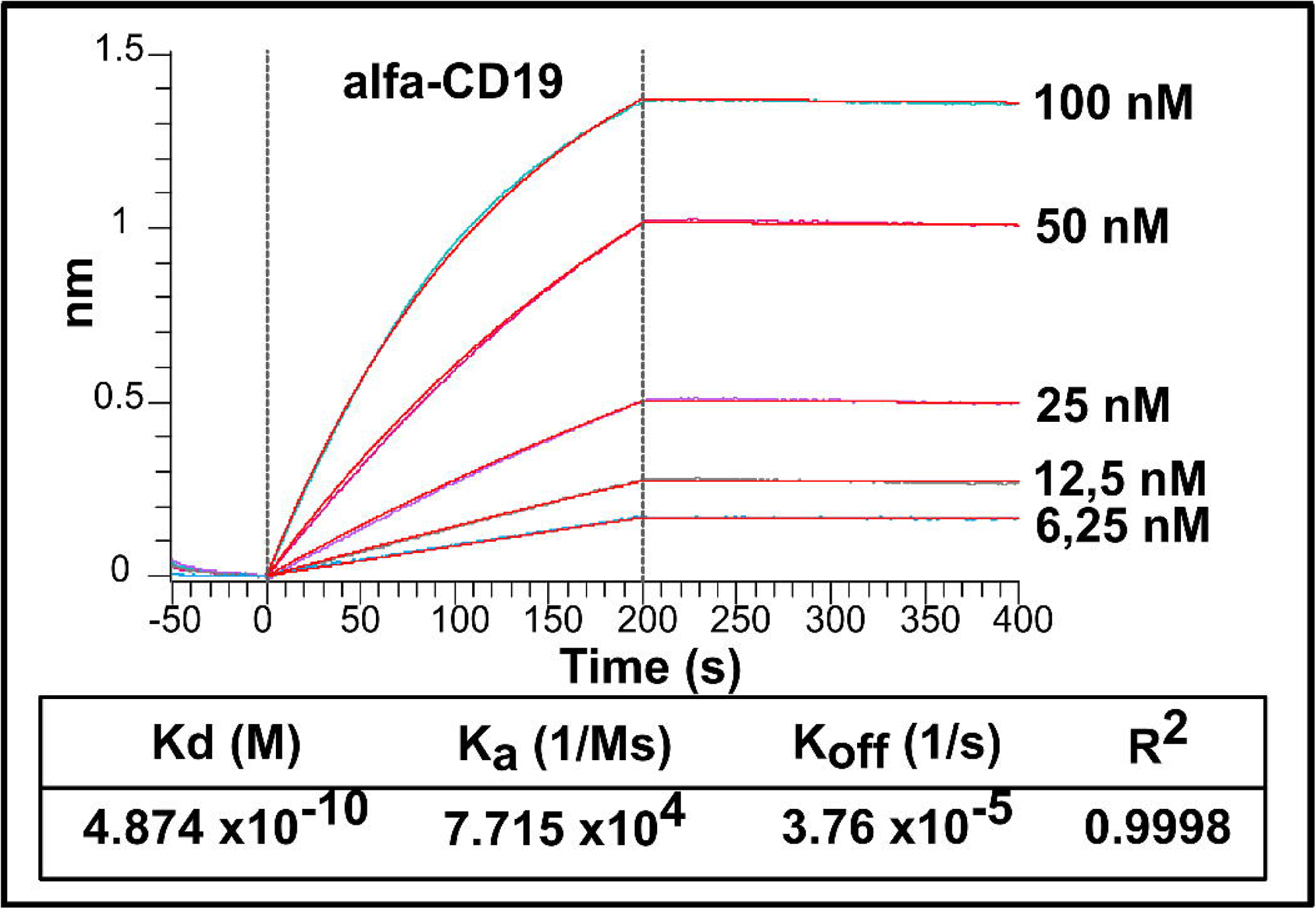
The kinetics of aNab:ALFA-tag interaction assessed by Octet R2 biosensor using a 1:1 binding model.

To express aNab on the cell surface two constructs were assembled (Figure 3a). These constructs consist of the aNab domain fused to a short flexible linker derived from the IgG hinge region in one case, and a long linker originated from a mutated IgG4 Fc fragment (mutation abrogates interaction with the Fc-receptor) in the other. Both constructs are flanked by IL-2 secretion signal and CD28 transmembrane domains from N- and C-termini respectively. They were cloned into the pWPXL vector (#12257, Addgene) under the control of the EF-1a promoter. Resulting vectors are referred to as pWPXL-aNab-Short and pWPXL-aNab-Long.

**Figure 3.**
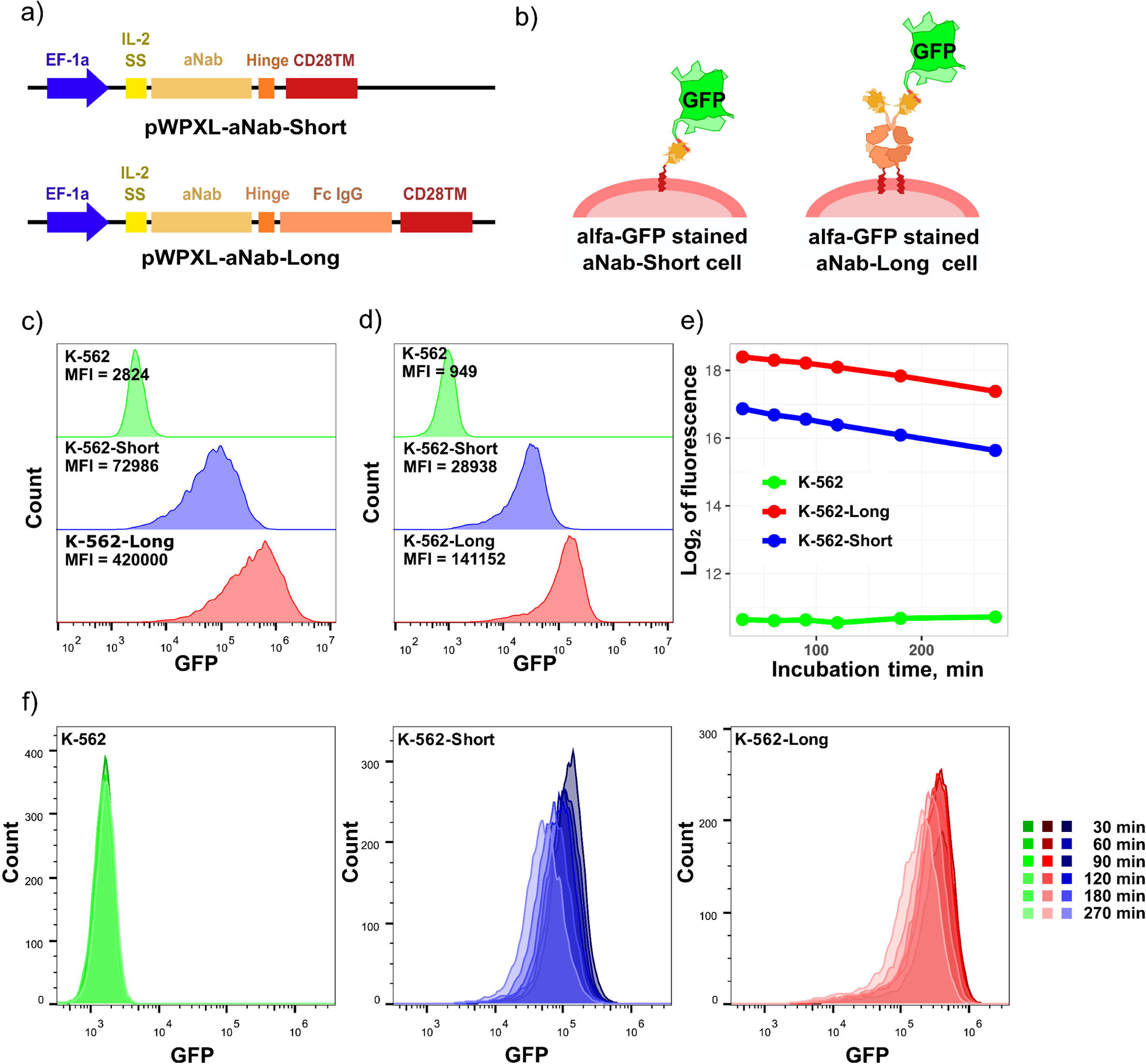
Schematic representation of pWPXL-aNab-Short and pWPXL-aNab-Long vectors (a); alfa-GFP stained cells (b); Flow cytometry analysis of transduced K-562 cells (c) before and (d) after selection step with alfa-GFP staining; The dynamics of alfa-GFP signal during cultivation of K-562, K-562-Short, and K-562-Long cells according to the flow cytometry (f); The dynamics of MFI values during incubation for K-562-Long or K-562-Short cells (e).

In order to perform functional control of aNab surface expression the ALFA-tagged GFP has been chosen (Figure 3b). A prokaryotic expression vector containing the alfa-GFP gene was designed, incorporating a His-tag for protein isolation, the GFP gene from pTurbo plasmid (#FP512, Eurogene), and the C-terminal ALFA-tag sequence (translated into PSRLEEELRRRLTE). This expression cassette was inserted into the pET28a vector (#69864-3, Addgene) for IPTG-inducible alfa-GFP expression in *E. coli* BL21 cells. The resulting alfa-GFP from cell lysate was purified using a Ni-NTA column with the Bio-Rad NGC Discovery system. The final yield was 70 mg of protein from 1 L of culture medium. The purity and quality of alfa-GFP were assessed via SDS-PAGE followed by Coomassie staining (Supplementary Figure 1b).

### 2.2 Lentiviral transduction and selection of K-562

In order to obtain stably transduced K-562 cell lines with the characterized level of aNab surface expression the lentiviral transduction was chosen. Lentiviral supernatants were produced using the pWPXL-aNab-Short and pWPXL-aNab-Long vectors in HEK293T cells. Subsequently, these supernatants were applied to the K-562 cells, and the transduction efficiency was assessed using alfa-GFP (Figure 3c). The pWPXL-aNab-Short and pWPXL-aNab-Long transduced cell lines are referred to as K-562-Short and K-562-Long, respectively. Both cell lines exhibited robust GFP signals, although with a differing modal expression levels as indicated by the Modal Fluorescence Intensity (MFI) values. Specifically, the normalized MFIs were determined to be 1:26:149 for K-562, K-562-Short, and K-562-Long, respectively.

Clone selection was conducted using a standard technique involving the dilution of cell cultures to stabilize the aNab levels on the cell surface. The presence and level of aNab expression were assessed via flow cytometry using recombinant alpha-GFP as the detection reagent. Selected clones of K-562-Long and K-562-Short and their GFP signals are presented in Figure 3d. Their MFIs are related as 1:30:149 for K-562, K-562-Short, and K-562-Long respectively.

### 2.3 Dynamics of immobilized alfa-GFP

Next, the dynamics of immobilized protein loss was assessed. Because of the viability of cells used as targets for the effector cells, the immobilization of any protein will be transient. That phenomenon is caused by the continuous expression of new aNab with subsequent membrane turnover. To access the turnover rate, the alfa-GFP staining was used. A series of samples were stained according to standard protocol, with incubation at 37°C, 5% CO2. At selected time points, samples from the series were analyzed using flow cytometry (Figure 3f). Given the linear relationship observed in the Log2(MFI) values (Figure 3e), it was possible to model the dynamics of GFP using a linear regression approach:

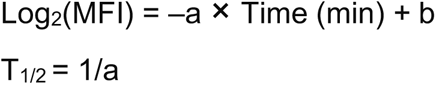

The half-life of alfa-GFP on the cell surface can be determined using linear regression models by calculating the division of 1 by the a-value (Supplementary 2), thereby yielding the time required for a 2X decrease in signal (e.g., MFI). The half-life for alfa-GFP is 196 minutes (3 hours and 16 minutes) for K-562-Short cells and 235 minutes (3 hours and 55 minutes) for K-562-Long cells. It is noteworthy that such calculations for K-562 cells cannot be performed due to a statistically insignificant a-value in this case (p-value = 0.182) and because its signal is assumed to be constant.

### 2.4 Immobilization of target protein on K-562-aNab-Short/Long cells

To validate this system, we chose CD19 protein as a target and FMC63 (Nicholson et al., 1997) as an antibody fragment in CAR because CD19 and FMC63 is the most described antibody-antigen pair in the entire CAR-T field. The ALFA-tagged CD19 protein was expressed and utilized for staining the model cell lines. The presence of alfa-CD19 on the cell surface was evaluated using a recombinant anti-CD19 FMC63 antibody. The CD19 knock-in K-562 cells are used as a positive control. The staining protocol comprised a 20-minute incubation with alfa-CD19 (similar to alfa-GFP), followed by washing and subsequent staining with the FMC63 antibody. It is noteworthy to say that the wild-type alfa-CD19 exhibited only a weak signal, as determined by Flow Cytometry using FMC63 antibodies (see Supplementary Figure 2a), which significantly decreased after purification and storage at 4°C (Supplementary Figure 2b). In contrast, the CD19 SF05 stability enhanced variant demonstrated a robust signal (Supplementary Figure 2c) in Flow Cytometry analysis with FMC63 antibodies. Both K-562-Short and K-562-Long cell lines exhibited membrane-bound CD19 levels comparable to those of CD19 knock-in cells.

### 2.5 Optimization of Reporter Assay Procedure

The ability to activate CAR signaling in CAR-T cells was initially evaluated using surrogate Jurkat-based reporter cells. Activation of CAR by surface antigen triggers the NFAT-response element in core promoter of GFP, allowing monitoring of antigen recognition via Flow Cytometry. When the Jurkat-based reporter cells are designated as Jurkat.NFAT cells, the Jurkat.NFAT.CAR cells are obtained by CAR introduction via lentiviral transduction. Both Jurkat.NFAT cells and Jurkat.NFAT.CAR cells were co-cultured with target cells according to the standard protocol with a series of 2-fold alfa-CD19 dilutions starting from 2 µg/ml of protein. After 24 hours of incubation, the cells were analyzed using Flow Cytometry. The percentage of activated cells was normalized by the proportion of CAR-positive cells (Figure 4a). The magnitude of activation was assessed as a relative value obtained by dividing the MFI of activated cells by that of resting cells. As expected, decreasing concentrations of alfa-CD19 led to reduced activation rates both in terms of the proportion of activated cells and the magnitude of activation (see Figure 4b). It is important to note that the concentration range of 0.5-2 µg/ml corresponds to a plateau of activation, indicating that this range is optimal for these experimental conditions.

**Figure 4.**
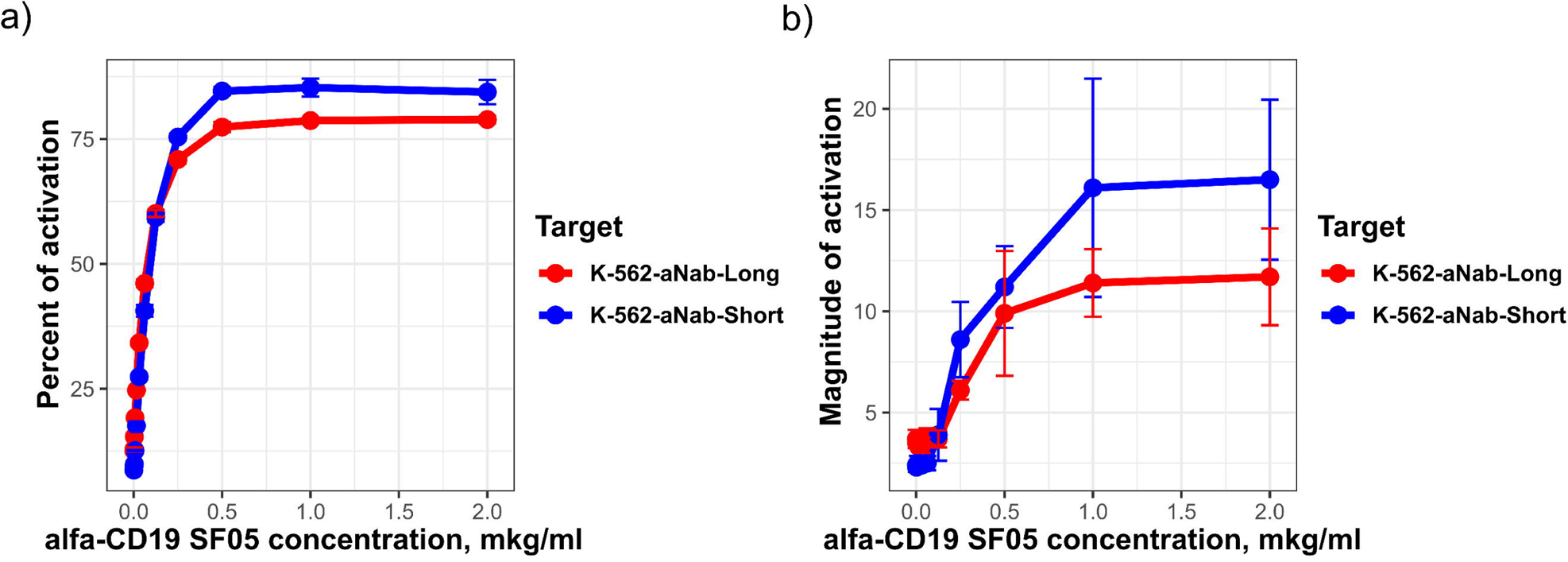
Titration of alfa-CD19 with NFAT reporters, the (a) percent of activation and (b) magnitude of activation compared to the negative population.

In certain instances, proteins themselves may induce CAR activation through receptor dimerization. This mechanism is commonly used to assess CAR activation in response to some antigens or other receptor clustering agents. To ensure that Jurkat.NFAT.CAR activation primarily arises from membrane-bound antigen and not from CAR crosslinking by soluble proteins, a series of control experiments was conducted. According to experiment results, a strong activation of reporter cells was only observed in the presence of antigen anchoring cells (K-562-Short and K-562-Long) together with CD19 in the medium, whereas soluble protein alone induced minimal 18% FMC63.CAR.NFAT reporter (FMC63.CAR) activation (Figure 5a).

**Figure 5.**
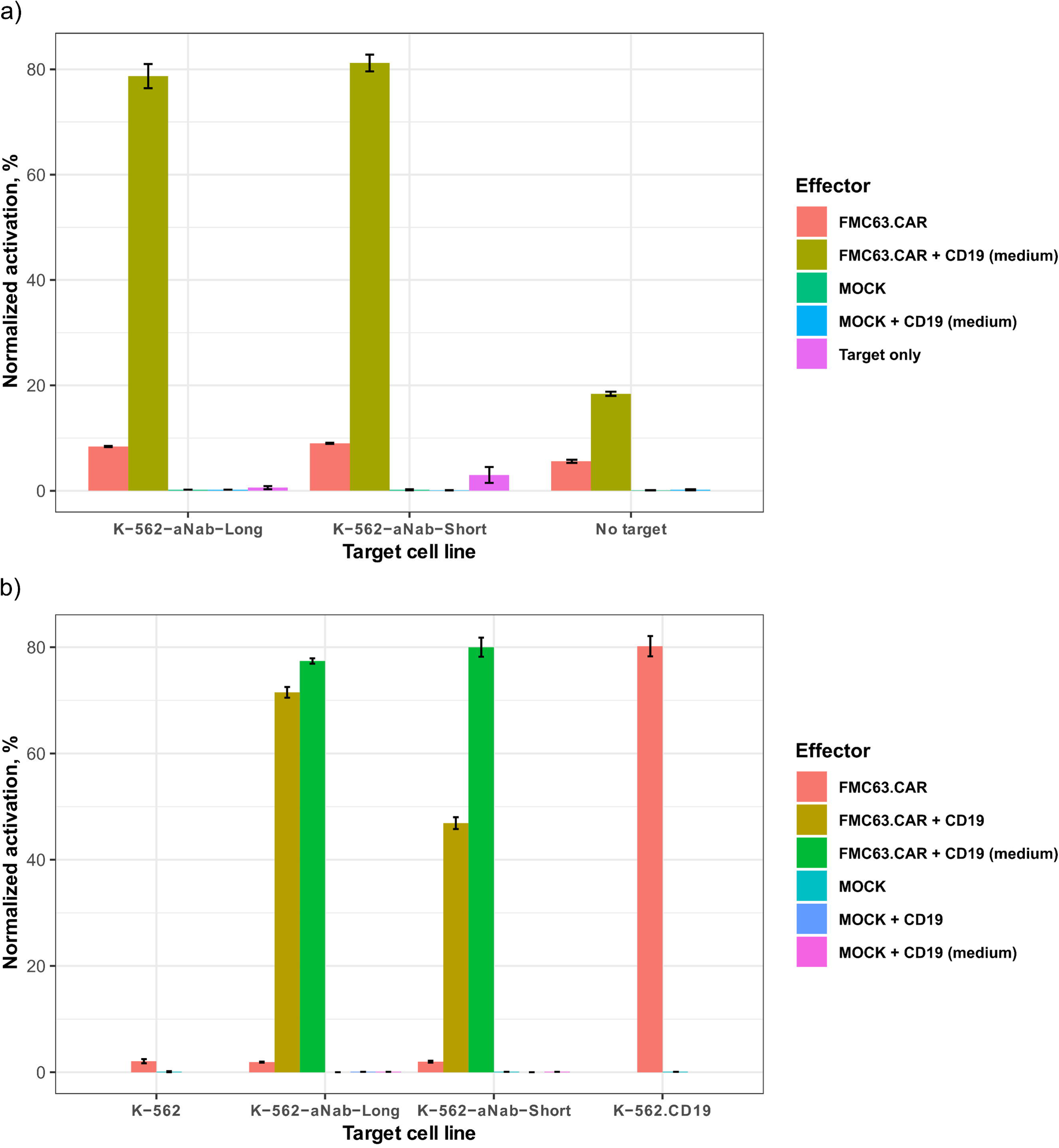
Evaluation of soluble CD19 mediated signaling in effector cells (a); The NFAT activation level in different experiment conditions and target cells (b).

To assess the difference in activation of anti-CD19 CAR by K-562-aNab-Short/Long, K-562 cells or K-562-CD19 cells a separate assay has been conducted (figure 5b). According to the experiment results, the strongest activation is observed in the case of K-562.CD19 cells. Interestingly, after model cell line staining before the experiment itself (FMC63 + CD19 samples), K-562-aNab-Short cells caused considerably reduced activation in Jurkat cells, but this effect fully reversed with the presence of the excess of soluble protein in the medium. In this case, ongoing staining with CD19 neutralizes the effect of membrane turnover to a degree, when the proportion of activated cells becomes almost the same as in positive control. Thus, *in situ* cell staining is shown to be the optimal experiment condition for this model.

## 3 Discussion

First, the ability of aNab to bind to its epitope was confirmed by Octet R2 assessment. This interaction exhibited a dissociation constant of approximately 0.5 nM (Figure 2) reaching sub-nanomolar range as in the original publication, validating the functional activity of the aNab:ALFA-tag pair.

Next, lentiviral transduction of K-562 cells was performed using aNab-Short and aNab-Long lentiviral vectors. In both cases, the majority of cells were aNab positive according to alfa-GFP staining, but K-562-Long cells exhibited approximately a 6-fold higher modal GFP fluorescence (Figure 3c). This expression pattern persisted even after selection (Figure 3d), and it can be explained by the difference in expression, transport to membrane, and membrane turnover.

The membrane turnover was assessed by staining selected cell lines with alfa-GFP, followed by a 4.5-hour incubation period with periodic Flow Cytometry measurements of remaining fluorescence. According to the Flow Cytometry data, GFP immobilization was shown to be transient (Figure 3e), with an approximate half-life of 196 min (3h 16m) for K-562-Short cells and 235 min (3h 55min) for K-562-Long cells. The difference in half-lives of immobilized proteins can contribute to the overall density of aNab on the membrane. This characteristic of the model system must be taken into consideration when conducting experiments with these cell lines as a model due to the limited time of antigen exposure by target cells.

While alfa-GFP protein was used to monitor the presence of aNab on the cell surface, alfa-CD19 protein was used as the target protein. CD19 protein is known to have low stability, resulting in aggregation, low expression yields, and loss of antigenic properties. The CD19 protein used here is a His- and ALFA-tagged protein (referred to as alfa-CD19). Initially, wild-type CD19 protein was expressed and purified for experimental purposes. The antigenic properties of this protein were assessed using specific anti-CD19 FMC63 antibodies, which also serve as antigen-recognizing parts of the CAR. Remarkably, only a weak signal caused by ALFA-tag immobilized CD19 was observed on the cells before purification, and an almost absent signal was observed after purification, size exclusion chromatography, followed by storage at 4°C (Supplementary Figure 2a and 2b). These results are consistent with our empirical experience with the use of wtCD19-based CAR detection kits, which lose their activity after short-term storage.

Due to the intrinsic instability of CD19 and the need to validate our model system, a mutated variant of CD19 was chosen. This variant, designated CD19 SF05 by Laurent and colleges (Laurent et al., 2021), bears three mutations within the compact locus (Immunoglobulin-fold 2) and lacks a fragment of the 5th exon, which encodes the peri-membrane part of the molecule. The main feature of this protein is a high degree of stability and non-impaired antigenic properties as assessed by FMC63 antibodies.

The amount of ALFA-tag immobilized CD19 was comparable to natively-expressed CD19 in the knock-in cell line (K-562.CD19 for short) (Supplementary Figure 2c). Importantly, such analysis revealed the non-impaired activity of alfa-CD19 (SF05) and its high degree of stability during storage, allowing desirable tests with this protein preparation.

The ability of K-562-Short and K-562-Long cells with immobilized protein to activate the CAR molecule was assessed using reporter Jurkat cells expressing GFP under the NFAT response element. The dynamic range of CD19 protein concentration was determined by co-culturing target cells with Jurkat.NFAT reporters. Increased concentration of CD19 protein resulted in a higher percentage of activation, with a plateau observed in the concentration range of 0.5-2 µg/ml (Figure 4a). Compared to the CD19 knock-in cell line, both K-562-Short and K-562-Long cells exhibited a similar percentage of activation of Jurkat.NFAT cells expressing the FMC63 CAR. While the percentage of activation is shown to be very consistent in our experimental condition (Figure 4a), the much more prominent variation in experimental outcome is shown for the magnitude of activation (Figure 4b), making this parameter less robust. Importantly, the transient nature of activation was particularly evident when cells were pre-stained before the test, resulting in reduced activation of reporter cells (Figure 5b). This emphasizes the importance of soluble protein utilization in this system for accurate results. Nonetheless, the activation of cells in the model cell lines (K-562-Short and K-562-Long cells) was comparable to that of the CD19 knock-in line by the percentage of activation (see Figure 5b). That shows the possibility of tagged protein anchoring on the cell surface using an anti-tag single-domain antibody (VHH) to produce antigen harboring model cell line for CAR-T cell evaluation.

## 4 Materials and methods

### 4.1 Cell lines and cultures

All bacterial strains were purchased from New England Biolabs (Ipswich, Massachusetts, USA), including *E. coli* strains NEB® 5-alpha and BL21. Bacterial cultures were supplemented with Luria-Bertani (LB) medium (#1231, Condalab, Spain).

Derived from the eukaryotic HEK293 cell line, FreeStyle™ 293-F cells (#R79007, Thermo Fisher, USA) utilized for construct expression were acquired from Thermo Fisher and cultured in FreeStyle™ 293 Expression Medium (#12338018, Thermo Fisher, USA) in accordance with the manufacturer’s recommendations.

The HEK 293T cells were obtained from the ATCC collection (USA) and supplemented with DMEM medium (#32430027, Gibco, USA) and 10% fetal bovine serum (FBS) (#16000044, Gibco, USA).

The K-562 cell line was obtained from the ATCC collection (USA) and maintained in RPMI medium (#11875093, Gibco, USA) supplemented with 10% FBS. Periodic screening for mycoplasma contamination in all eukaryotic cell lines was conducted via PCR using specific primers targeting mycoplasma 16S rRNA.

Jurkat.NFAT reporter cells were gracefully provided by Hanna Klych and cultured in the same matter as K-562 cells.

### 4.2 Vector assembly

Several vectors were designed in this study for the following purposes: 1) to demonstrate the interaction between aNab and the ALFA tag, 2) to produce ALFA-tagged proteins for aNab monitoring, and 3) to generate lentiviral plasmids for co-transfection and lentivirus production. The sequence of aNab was synthesized according to the original study (H. Götzke et al., 2019) by Synbio Technologies (Monmouth Junction, NJ, USA) in the form of a plasmid harboring aNab. Subsequently, the aNab gene was amplified using the 5’-CTACGGGGATCCACCATGTACAGGATGCAACTCC-3’ and 5’-GAATTATGCGGCCGCGCTGCTAACGGTAAC-3’ primers, followed by cloning into the pWPXL and pET20b_VHH vectors using NcoI and NotI restriction sites.

The pET20b_VHH_aNab vector was utilized to produce aNab fused with an IgA hinge fragment and AviTag. This recombinant protein was biotinylated for further assessment using the avidin-utilizing Octet R2 (Sartorius, Germany).

According to the concept, aNab can be anchored on the outer membrane of the target cell. Hence, the full construct comprises an IL-2 signal sequence, aNab, a short or long linker, and the CD28 transmembrane domain. The linkers utilized consist of the IgG hinge region for Linker_S or the IgG Fc domain with the IgG hinge for the Linker_L. The assembly of constructs was performed in the pWPXL lentiviral vector (#12257, Addgene) under the constant EF-1a promoter, followed by the components described previously. The resulting vectors, referred to as pWPXL-aNab-Short and pWPXL-aNab-Long, were used for co-transfection with lentiviral helper plasmids.

Due to the absence of any eukaryotic selection marker in pWPXL, the functional activity of aNab at the cell surface was chosen for monitoring. Therefore, ALFA-tagged GFP was produced to analyze the presence of aNab. The TurboGFP gene from pULTRA was cloned using a pair of primers 5’-CACGGATCCGGCATGGAGAGCGACGAGAGCGG and 5’-TTAGCGGCCGCTTCTTCACCGG. This resulted in a PCR product with two flanking restriction sites – NotI and BamHI. Simultaneously, the vector pET-28a (#69864-3, Addgene) was amplified using 5’-GAAGCGGCCGCCCCGTCTCGTCTGGAAGAAGAACTGCGTCGTCGTCTGACCGAATAACACCACCACCACCACCACCACTGAGATCCGGC and 5’-GCCGGATCCGTGATGATGATGATGATGGCTGCTGCCC to introduce NotI and BamHI restriction sites into the vector backbone, as well as the sequence of the ALFA-tag (subsequently translated to PSRLEEELRRRLTE). The resulting PCR products were digested with the corresponding restrictases and ligated to assemble the pET-28a_ALFA_GFP expression vector for prokaryotic expression.

### 4.3 Bacterial expression

E. coli BL21 cells were transformed with either the pET-28a_ALFA_GFP or pET20b_VHH_aNab plasmid DNA preparations, and subsequently utilized for bacterial expression. A 1 ml overnight culture was inoculated into 1 L of sterile Terrific Broth (#1246, Condalab, Spain) medium supplemented with the appropriate antibiotic (kanamycin at a final concentration of 50 μg/ml or ampicillin at a final concentration of 100 μg/ml) and incubated at 37°C with agitation until reaching an OD600 of approximately 1-1.2. Following this, induction was initiated by the addition of IPTG to a final concentration of 0.2 mM, and incubation continued for approximately 17 hours at 30°C with agitation at approximately 150 rpm.

Upon completion of expression, cells were harvested by centrifugation at 3000g, 4°C for 20 minutes, and subsequently homogenized using an Emulciflex C5 (Avestin inc., Ottawa, Canada). The resulting cell lysate underwent clarification by ultracentrifugation at 90,000 g, 4°C, for 50 minutes to remove cellular debris. The clarified solution was then applied to an IMAC column for further purification of alfa-GFP and aNab-AviTag. After purification and subsequent dialysis in PBS, the purified alfa-GFP and aNab-AviTag proteins were analyzed by SDS-PAGE and Coomassie staining (Supplementary Figure 1a and 1b). The remaining protein solution was supplemented with 10% glycerol and stored at −70°C.

### 4.4 Eukaryotic Expression

Plasmid DNA was purified using the PureLink™ HiPure Plasmid Miniprep Kit (#K210003, Thermo Fisher, USA) for eukaryotic expression in the FreeStyle™ 293 cell line. A total of 30 × 10^6 cells in 30 ml of FreeStyle™ 293 Expression Medium (#12338026, Thermo Fisher, USA) were transfected with 30 µg of DNA. The plasmid transfection procedure involved diluting 30 µg of purified plasmid DNA in 750 µl of Opti-MEM™ (#11058021, Thermo Fisher, USA), along with dilution of 90 µg of PEI in 750 µl of Opti-MEM™ (DNA:PEI ratio of 1:3 by mass). Subsequently, the PEI solution in Opti-MEM™ was added to the DNA solution, mixed thoroughly, and incubated for 10 minutes at room temperature. The resulting solution was then added to the FreeStyle™ 293 cells, followed by a 5-day incubation at 37°C with 5% CO2 under constant agitation in a humidified atmosphere. The culture medium was clarified by sequential centrifugation at 600 × g for cell sedimentation and 90,000 × g for maximal clean-up. This clarified medium was then utilized for protein purification.

### 4.5 CD19 Mutagenesis

Due to the notable instability of wild-type (WT) CD19, a stabilized form of this protein was utilized for system verification. This form harbors three point mutations (M75V, R76S, and F85S) as well as an additional truncation of 13 residues from exon 5 (Laurent et al., 2021). The CD19 gene was cloned into the pFUSE vector using the primers 5’-CACGAATTCGGCCCCCGAGGAACCTCTAGTG and 5’-TCTGCGGCCGCCTTCCAGCCACCAGTCCTCAG and the EcoRI and NotI restriction enzymes. Additionally, His and ALFA tags were added to the C-terminus of the protein. The resulting protein is referred to as alfa-CD19wt. Mutagenesis was performed through two sequential amplifications with primer pair 1 (5’-CCAGGGGCCTCATGTGGATTCCC and 5’-CCATCTGGCTTTTCATCTCCAACGTCTCTCAACAGATGGG) and primer pair 2 (5’-GTGGATTCCCAGGCCTGGCAGCCC and 5’-GTGAGCCCCCTGGCCATCTGGCTTTTC). The amplification scheme included sequential reactions with blunt-end ligation after each step. The resulting gene includes the mutated CD19 and His and ALFA tags, and it is referred to as alfa-CD19.

### 4.6 Ni-NTA Chromatography

Ni-NTA chromatography was conducted using the Bio-Rad NGC Chromatography System with either a 1- or 5-ml Ni-NTA column. *E. coli* lysate was applied to the pre-equilibrated column and subsequently washed with high-saline PBS supplemented with 25 mM imidazole. Elution of the bound protein was achieved by a gradual increase in imidazole concentration from 25 to 250 mM. Fractions exhibiting peaked concentrations of protein (based on OD280) were collected for further dialysis in PBS.

CD19 isolation by Ni-NTA chromatography followed by size exclusion chromatography was used to get rid of any possible aggregates in purified protein sample. The resulting protein was stored in PBS with 300mM NaCl.

### 4.7 Flow Cytometry

Flow Cytometry analysis was performed using alfa-GFP, a derivative of turboGFP. Cell staining with alfa-GFP was carried out at a final concentration of 1 µg/ml for 30 minutes, followed by a double wash with PBS to remove unbound alfa-GFP. Measurements were performed using the Beckman Coulter CytoFLEX with a 488 nm laser and a 525/40 bandpass filter to analyze the presence of GFP. Detection of alfa-CD19 was performed using FLAG-tagged FMC63 antibody with anti-FLAG secondary antibodies (IC8529R, R&D Systems, USA). The measurement process involved the APC channel (648 nm laser with a 660/10 BP filter). Data analysis and visualization were performed using FlowJo software version 10.0.07.

### 4.8 Production of Lentiviruses

The production of lentiviruses involved co-transfection of the lentiviral plasmid vector system. This system included a pWPXL transfer vector (Addgene, #12257) with functional long terminal repeats (LTRs) and *ψ* site for packaging, a pCMV-dr8_91 (Addgene, #2221) packaging vector with polyprotein necessary for lentiviral particle production, and a pMD2.G (Addgene, #12259) vector with vesicular stomatitis glycoprotein (VSV-G) for pseudotyping. HEK 293T cells were plated in 100 mm Petri dishes and grown to approximately 50% confluence on the day of transfection. The culture medium was changed to fresh serum-free DMEM on the day of transfection. The co-transfection procedure was similar to the previously described transfection but required 10 µg of pWPXL, 6 µg of pCMVdr8_91, and 3 µg of pMD2.G plasmids. After 16 hours of incubation, the medium was replaced with fresh DMEM supplemented with 10% FBS. Lentivirus-containing supernatant was collected after 48 hours of expression and underwent syringe filtering with 0.45 µm PET filter to remove any cells or debris, followed by centrifugation at 2,000g for 16 hours to concentrate the viruses for transduction. After disposing of the supernatant, the residual medium (∼100 µl) was used to resuspend the lentivirus-containing pellet. This solution was aliquoted and stored at −70°C.

### 4.9 Lentiviral Transduction

Concentrated supernatants were used to transduce K-562 cells. A total of 30,000 cells per well were plated in 96-well plates, followed by the addition of concentrated supernatant (100, 50, 25, and 12 µl per well) in a final volume of 0.2 ml. Additionally, 1 µg of Polybrene (TR-1003, Sigma-Aldrich, USA) per well was added to enhance transduction efficiency. The cells were mixed using a pipette and then spinoculated by centrifugation at 500g for 1.5 hours at 37°C. Spinoculated cells were subsequently cultured in a humidified incubator at 37°C with 5% CO2.

After transduction procedure the modificated cells were selected by serial dilutions in 96-well plates. Cell lines from the most diluted wells were further analyzed to asses clone homogeneity by alfa-GFP staining and Flow Cytometry.

### 4.10 NFAT Reporter Assay

A GFP under the control of NFAT response element in Jurkat cells was utilized to assess the activation of FMC63 CAR in cells. The Jurkat cells with the NFAT reporter system were transduced with a LV particles encoding CAR construct containing FMC63scFv + CD28TM + 4-1BB + CD3z with EGFRt as a selective marker. These cells were enriched in a selective column with EGFRt-binding antibodies and used as a positive effector T-cells. The Jurkat cells with the NFAT reporter system but without CAR transduction were used as negative effector T-cells. The test procedure involved 24 hours of co-culturing effectors (3 × 10^4 per well) with target cells (1 × 10^4 per well), followed by Flow Cytometry analysis in the FITC channel (BP 525/40).

### 4.11 NFAT Reporter Assay Interpretation

Data analysis and interpretation involved the gating of Jurkat cells by FSC/SSC, followed by the evaluation of the proportion of GFP-positive activated cells. Further analysis required the normalization of activated Jurkat cells to the proportion of CAR+ cells in the enriched population. The magnitude of cell activation was assessed by calculating the Modal Fluorescence intensity (MFI) at the FITC channel for activated and resting cells, followed by the division of the MFI of activated cells by the MFI of resting cells. This number represented the X-fold increase in signal compared to the background.

---

**Supplementary Figure 1.**
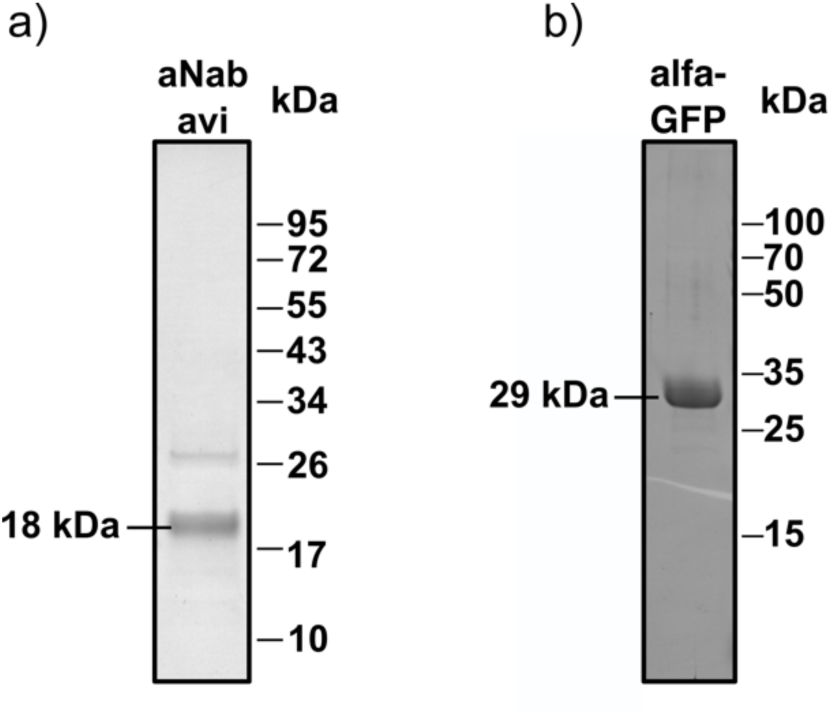
The electrophoretic analysis of (a) aNab-avi and (b) alfa-GFP via SDS-PAGE followed by Coomassie staining

**Supplementary 2**

The equations of GFP dynamics:

Log_2_(MFI) = – a*Time (min) + b

Thus, the increase of Log_2_(MFI) by 1 means 2-fold increase in fluorescence so it can be coupled with time interval half-life for alfa-GFP (T_1/2_). We consider that at the time interval Time_2_ – Time_1_ = T_1/2_ and the Log_2_(MFI) value decreases by 1:

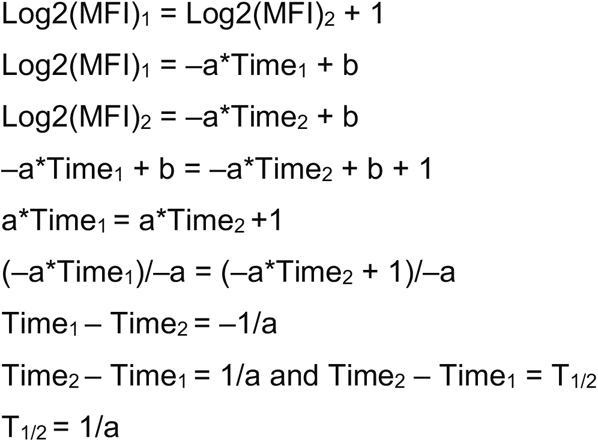

The a-values and b-values for all samples from linear regression models in R (version 3.6.3) are presented in Table 2.

**Table 2.**
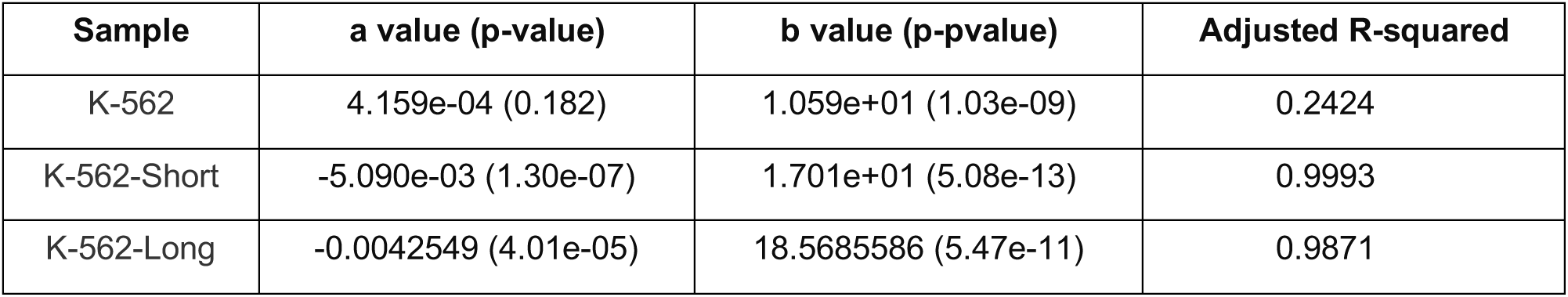
The statistical parameters of linear regression components.

**Supplementary Figure 2.**
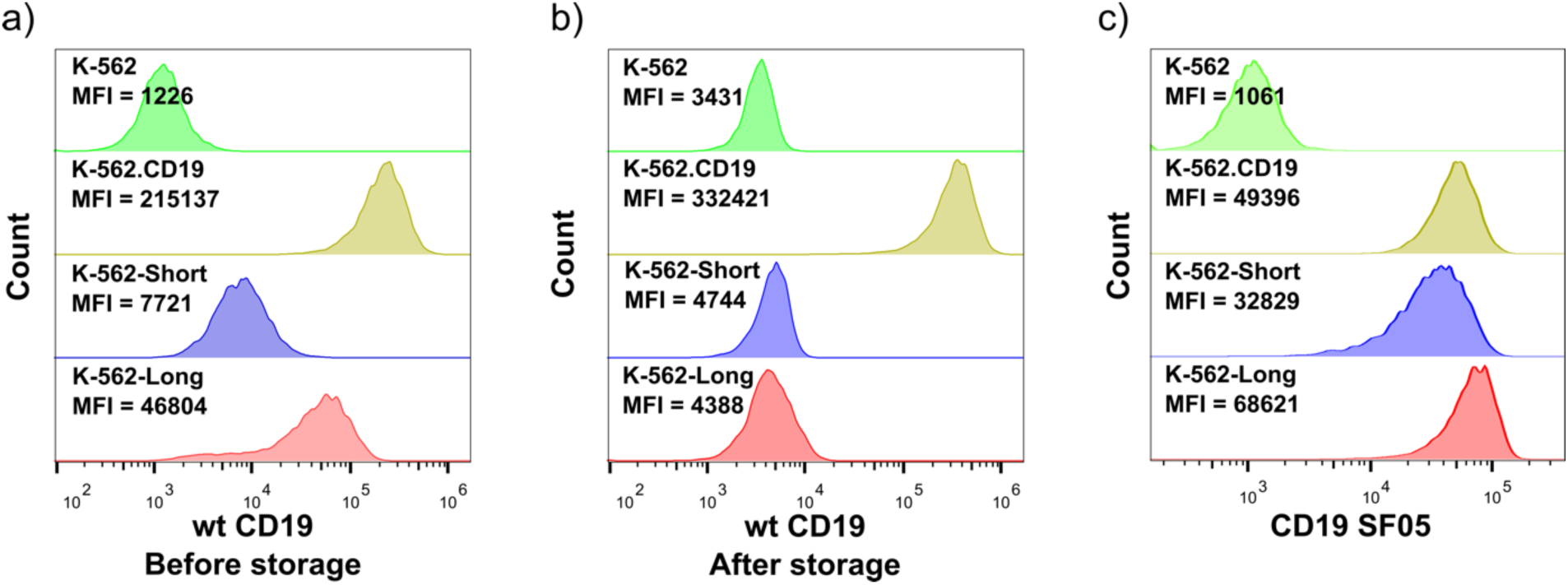
Comparation of wt CD19 before purification and storage (a) and after (b) on the cell surface of model cell lines as assessed by FMC63 antibodies to K-562 cells and CD19 knock-in K-562 cells. The same analysis of purified CD19 SF05 (c).

## Acknowledgments

The authors are highly grateful to Hanna Klych for providing Jurkat NFAT reporter cells, and to Michail Shapira for protein isolation and size exclusion chromatography.

## References

Davenport, A.J., Cross, R.S., Watson, K.A., Liao, Y., Shi, W., Prince, H.M., Beavis, P.A., Trapani, J.A., Kershaw, M.H., Ritchie, D.S., Darcy, P.K., Neeson, P.J., Jenkins, M.R., 2018. Chimeric antigen receptor T cells form nonclassical and potent immune synapses driving rapid cytotoxicity. Proc. Natl. Acad. Sci. U. S. A. 115, E2068–E2076. 10.1073/pnas.1716266115

Fujiwara, K., Tsunei, A., Kusabuka, H., Ogaki, E., Tachibana, M., Okada, N., 2020. Hinge and Transmembrane Domains of Chimeric Antigen Receptor Regulate Receptor Expression and Signaling Threshold. Cells 9, 1182. 10.3390/cells9051182

Götzke, H., Kilisch, M., Martínez-Carranza, M., Sograte-Idrissi, S., Rajavel, A., Schlichthaerle, T., Engels, N., Jungmann, R., Stenmark, P., Opazo, F., Frey, S., 2019. The ALFA-tag is a highly versatile tool for nanobody-based bioscience applications. Nat. Commun. 10, 4403. 10.1038/s41467-019-12301-7

Haso, W., Lee, D.W., Shah, N.N., Stetler-Stevenson, M., Yuan, C.M., Pastan, I.H., Dimitrov, D.S., Morgan, R.A., FitzGerald, D.J., Barrett, D.M., Wayne, A.S., Mackall, C.L., Orentas, R.J., 2013. Anti-CD22-chimeric antigen receptors targeting B-cell precursor acute lymphoblastic leukemia. Blood 121, 1165–1174. 10.1182/blood-2012-06-438002

Huang, J., Brameshuber, M., Zeng, X., Xie, J., Li, Q., Chien, Y., Valitutti, S., Davis, M.M., 2013. A Single Peptide-Major Histocompatibility Complex Ligand Triggers Digital Cytokine Secretion in CD4+ T Cells. Immunity 39, 846–857. 10.1016/j.immuni.2013.08.036

Irvine, D.J., Purbhoo, M.A., Krogsgaard, M., Davis, M.M., 2002. Direct observation of ligand recognition by T cells. Nature 419, 845–849. 10.1038/nature01076

Kantari-Mimoun, C., Barrin, S., Vimeux, L., Haghiri, S., Gervais, C., Joaquina, S., Mittelstaet, J., Mockel-Tenbrinck, N., Kinkhabwala, A., Damotte, D., Lupo, A., Sibony, M., Alifano, M., Dondi, E., Bercovici, N., Trautmann, A., Kaiser, A.D., Donnadieu, E., 2021. CAR T-cell Entry into Tumor Islets Is a Two-Step Process Dependent on IFNγ and ICAM-1. Cancer Immunol. Res. 9, 1425–1438. 10.1158/2326-6066.CIR-20-0837

Laurent, E., Sieber, A., Salzer, B., Wachernig, A., Seigner, J., Lehner, M., Geyeregger, R., Kratzer, B., Jäger, U., Kunert, R., Pickl, W.F., Traxlmayr, M.W., 2021. Directed Evolution of Stabilized Monomeric CD19 for Monovalent CAR Interaction Studies and Monitoring of CAR-T Cell Patients. ACS Synth. Biol. 10, 1184–1198. 10.1021/acssynbio.1c00010

Lei, W., Zhao, A., Liu, H., Yang, C., Wei, C., Guo, S., Chen, Z., Guo, Q., Li, L., Zhao, M., Wu, G., Ouyang, G., Liu, M., Zhang, J., Gao, J., Qian, W., 2024. Safety and feasibility of anti-CD19 CAR T cells expressing inducible IL-7 and CCL19 in patients with relapsed or refractory large B-cell lymphoma. Cell Discov. 10, 1–14. 10.1038/s41421-023-00625-0

Liu, Y., Xu, X., Liu, D., Wu, X., Gao, Y., Wang, H., Yan, F., Yang, W., Zhao, D., He, F., Tang, L., 2023. 30-color full spectrum flow cytometry panel for deep immunophenotyping of T cell subsets in murine tumor tissue. J. Immunol. Methods 516, 113459. 10.1016/j.jim.2023.113459

Malik-Chaudhry, H.K., Prabhakar, K., Ugamraj, H.S., Boudreau, A.A., Buelow, B., Dang, K., Davison, L.M., Harris, K.E., Jorgensen, B., Ogana, H., Pham, D., Schellenberger, U., Van Schooten, W., Buelow, R., Iyer, S., Trinklein, N.D., Rangaswamy, U.S., n.d. TNB-486 induces potent tumor cell cytotoxicity coupled with low cytokine release in preclinical models of B-NHL. mAbs 13, 1890411. 10.1080/19420862.2021.1890411

Mukherjee, M., Mace, E.M., Carisey, A.F., Ahmed, N., Orange, J.S., 2017. Quantitative Imaging Approaches to Study the CAR Immunological Synapse. Mol. Ther. 25, 1757–1768. 10.1016/j.ymthe.2017.06.003

Nerreter, T., Letschert, S., Götz, R., Doose, S., Danhof, S., Einsele, H., Sauer, M., Hudecek, M., 2019. Super-resolution microscopy reveals ultra-low CD19 expression on myeloma cells that triggers elimination by CD19 CAR-T. Nat. Commun. 10, 3137. 10.1038/s41467-019-10948-w

Nicholson, I.C., Lenton, K.A., Little, D.J., Decorso, T., Lee, F.T., Scott, A.M., Zola, H., Hohmann, A.W., 1997. Construction and characterisation of a functional CD19 specific single chain Fv fragment for immunotherapy of B lineage leukaemia and lymphoma. Mol. Immunol. 34, 1157–1165. 10.1016/s0161-5890(97)00144-2

Palomba, M.L., Qualls, D., Monette, S., Sethi, S., Dogan, A., Roshal, M., Senechal, B., Wang, X., Rivière, I., Sadelain, M., Brentjens, R.J., Park, J.H., Smith, E.L., 2022. CD19-directed chimeric antigen receptor T cell therapy in Waldenström macroglobulinemia: a preclinical model and initial clinical experience. J. Immunother. Cancer 10, e004128. 10.1136/jitc-2021-004128

Pandita, A., Aldape, K.D., Zadeh, G., Guha, A., James, C.D., 2004. Contrasting in vivo and in vitro fates of glioblastoma cell subpopulations with amplified EGFR. Genes. Chromosomes Cancer 39, 29–36. 10.1002/gcc.10300

Schuster, S.J., Zhang, J., Yang, H., Agarwal, A., Tang, W., Martinez-Prieto, M., Bollu, V., Kuzan, D., Maziarz, R.T., Kersten, M.J., 2022. Comparative efficacy of tisagenlecleucel and lisocabtagene maraleucel among adults with relapsed/refractory large B-cell lymphomas: an indirect treatment comparison. Leuk. Lymphoma 63, 845–854. 10.1080/10428194.2021.2010069

Si, X., Xiao, L., Brown, C.E., Wang, D., 2022. Preclinical Evaluation of CAR T Cell Function: In Vitro and In Vivo Models. Int. J. Mol. Sci. 23, 3154. 10.3390/ijms23063154

Uhlén, M., Fagerberg, L., Hallström, B.M., Lindskog, C., Oksvold, P., Mardinoglu, A., Sivertsson, Å., Kampf, C., Sjöstedt, E., Asplund, A., Olsson, I., Edlund, K., Lundberg, E., Navani, S., Szigyarto, C.A.-K., Odeberg, J., Djureinovic, D., Takanen, J.O., Hober, S., Alm, T., Edqvist, P.-H., Berling, H., Tegel, H., Mulder, J., Rockberg, J., Nilsson, P., Schwenk, J.M., Hamsten, M., von Feilitzen, K., Forsberg, M., Persson, L., Johansson, F., Zwahlen, M., von Heijne, G., Nielsen, J., Pontén, F., 2015. Proteomics. Tissue-based map of the human proteome. Science 347, 1260419. 10.1126/science.1260419

Zhang, Y., Patel, R.P., Kim, K.H., Cho, H., Jo, J.-C., Jeong, S.H., Oh, S.Y., Choi, Y.S., Kim, S.H., Lee, J.H., Angelos, M., Guruprasad, P., Cohen, I., Ugwuanyi, O., Lee, Y.G., Pajarillo, R., Cho, J.H., Carturan, A., Paruzzo, L., Ghilardi, G., Wang, M., Kim, S., Kim, S.-M., Lee, H.-J., Park, J.-H., Cui, L., Lee, T.B., Hwang, I.-S., Lee, Y.-H., Lee, Y.-J., Porazzi, P., Liu, D., Lee, Y., Kim, J.-H., Lee, J.-S., Yoon, D.H., Chung, J., Ruella, M., 2023. Safety and efficacy of a novel anti-CD19 chimeric antigen receptor T cell product targeting a membrane-proximal domain of CD19 with fast on- and off-rates against non-Hodgkin lymphoma: a first-in-human study. Mol. Cancer 22, 200. 10.1186/s12943-023-01886-9

