## Supplementary 2 for "Modular Cell Line for Scalable and Rapid In Vitro Evaluation of Chimeric Antigen Receptors"

The equations of GFP dynamics:

Log_2_(MFI) = – a*Time (min) + b

Thus, the increase of Log_2_(MFI) by 1 means 2-fold increase in fluorescence so it can be coupled with time interval half-life for alfa-GFP (T_1/2_). We consider that at the time interval Time_2_ – Time_1_ = T_1/2_ and the Log_2_(MFI) value decreases by 1:

Log2(MFI)_1_ = Log2(MFI)_2_ + 1

Log2(MFI)_1_ = –a*Time_1_ + b

Log2(MFI)_2_ = –a*Time_2_ + b

–a*Time_1_ + b = –a*Time_2_ + b + 1

a*Time_1_ = a*Time_2_ +1

(–a*Time_1_)/–a = (–a*Time_2_ + 1)/–a

Time_1_ – Time_2_ = –1/a

Time_2_ – Time_1_ = 1/a and Time_2_ – Time_1_ = T_1/2_

T_1/2_ = 1/a

The a-values and b-values for all samples from linear regression models in R (version 3.6.3) are presented in Table 2.

Table 2 The statistical parameters of linear regression components.

| **Sample** | **a value (p-value)** | **b value (p-pvalue)** | **Adjusted R-squared** |
| --- | --- | --- | --- |
| K-562 | 4.159e-04 (0.182) | 1.059e+01 (1.03e-09) | 0.2424 |
| K-562-Short | -5.090e-03 (1.30e-07) | 1.701e+01 (5.08e-13) | 0.9993 |
| K-562-Long | -0.0042549 (4.01e-05) | 18.5685586 (5.47e-11) | 0.9871 |
