## Supplementary Figure 1 for "Modular Cell Line for Scalable and Rapid In Vitro Evaluation of Chimeric Antigen Receptors"

The electrophoretic analysis of (a) aNab-avi and (b) alfa-GFP via SDS-PAGE followed by Coomassie staining

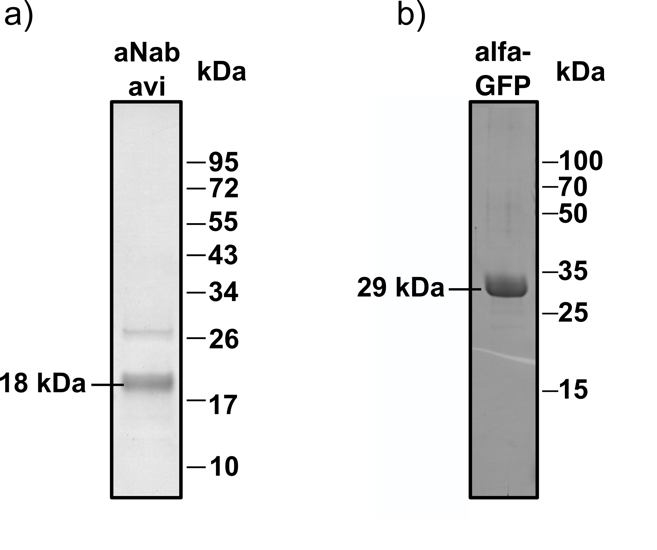
