## Supplementary Figure 2 for "Modular Cell Line for Scalable and Rapid In Vitro Evaluation of Chimeric Antigen Receptors"

Comparation of wt CD19 before purification and storage (a) and after (b) on the cell surface of model cell lines as assessed by FMC63 antibodies to K-562 cells and CD19 kno
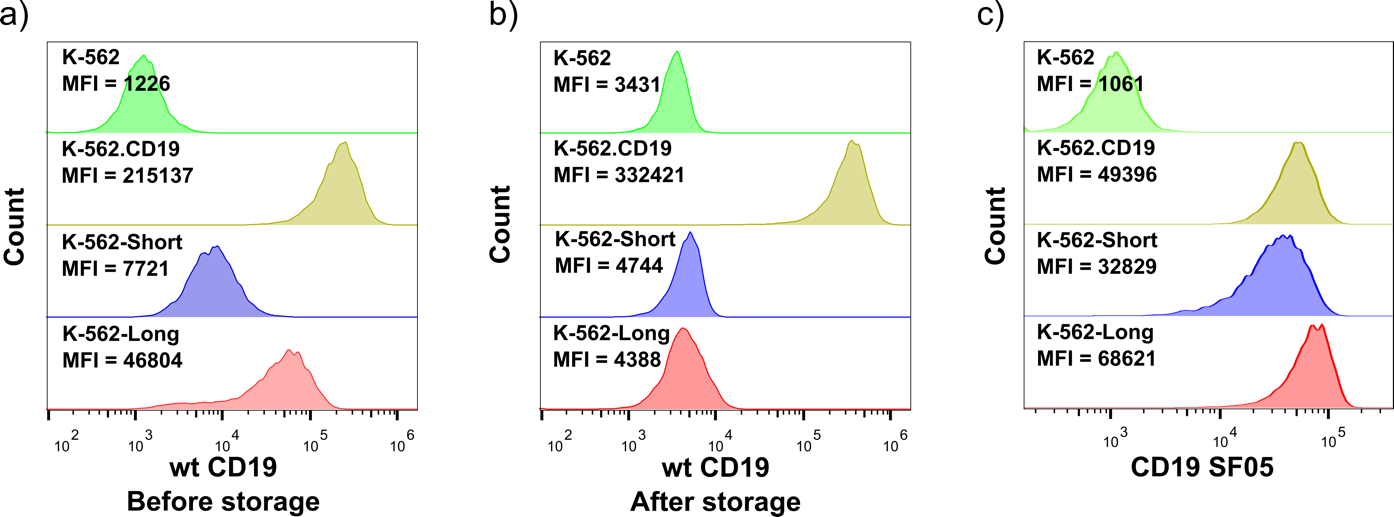
ck-in K-562 cells. The same analysis of purified CD19 SF05 (c).
